# Phenotype and functionality of follicular helper T cells in patients with acute dengue infection

**DOI:** 10.1101/723460

**Authors:** Ayesha M. Wijesinghe, Jayani Gamage, Hemantha Goonewardana, Laksiri Gomes, Deshni Jayathilaka, Dulharie T. Wijeratne, Ruklanthi de Alwis, Chandima Jeewandara, Ananda Wijewickrama, Graham S. Ogg, Gathsaurie Neelika Malavige

**Affiliations:** Centre for Dengue Research, University of Sri Jayewardenepura, Nugegoda, Sri Lanka; National Institute of Infectious Diseases, Angoda, Sri Lanka; MRC Human Immunology Unit, Weatherall Institute of Molecular Medicine, University of Oxford, Oxford, United Kingdom; Programme in Emerging Infectious Diseases, Duke-NUS Medical School, Singapore; Viral Research & Experimental Medicine Centre, SingHealth/Duke-NUS, Singapore

## Abstract

**Background:** The association of functionality and phenotype of follicular helper T cells (Tfh) with dengue virus (DENV) specific antibody responses and clinical disease severity is has not been well studied.

**Methodology/Principal findings:** We investigated the phenotype and functionality of Tfhs in adult patients (DF = 18, DHF = 22) with acute dengue of varying severity using multiparametric flowcytometry. We determined if the properties of Tfhs were correlated with viraemia, disease severity, plasmablast responses and DENV-specific serum antibody responses. We further evaluated the changes in neutralizing antibodies (Neut50) with viraemia and clinical disease severity in a different cohort of patients with acute secondary DENV1 (DF=12, DHF=10) and DENV2 infection (DF=8, DHF=9).

Tfhs (especially those producing IL-21 and co-expressing PD-1 and ICOS) were found to be significantly expanded (p<0.0001) and highly activated in patients with DHF compared to those with DF. The frequency of Tfh cells significantly correlated with DENV-specific IgG, NS1-specific antibodies and Neut50 antibody titres, which were also significantly higher in patients with DHF. Although the Neut50 titres increased during the course of acute secondary DENV infection, they showed differences based on serotype. For instance, the Neut50 titres were significantly higher during the latter part of illness in patients with DF compared DHF in DENV1 infection, while in DENV2, patients with DHF had significantly higher titres. The viral loads during early illness did not correlate with the subsequent rise in the Neut50 antibody titres during time point of illness.

**Conclusions/Significance:** The expansion of Tfhs is associated with DHF and DENV-specific IgG, NS1-specific and neutralising antibodies. Neut50 titres did not associate with disease severity or viraemia at the point of first presentation during the febrile phase, but later titres do show differential association with severity in patients with DENV1 compared to DENV2.

**Author summary:** Follicular helper T cells (Tfh cells) are a subset of T cells which are important in activation of germinal centre T cells and induction of long-lasting virus specific antibodies. The association of the functional status and phenotype of Tfh cells in relation to dengue virus (DENV) specific antibodies, clinical disease severity and the degree of viraemia in patients with acute dengue has not been extensively studied. Here we show that the Tfh cells are significantly expanded in patients with severe clinical disease and associate with levels of serum DENV envelop specific IgG, NS1-specific antibodies and neutralising antibodies (Neu50). Although DENV Neu50 antibody titres are thought to associate with protection, the Neut50 antibody titres were similar in patients who proceeded to develop mild or severe clinical disease and also the increase in their titres showed significant variation based on infecting DENV serotype and clinical disease severity. Therefore, although Tfh cells are expanded in patients with more severe forms of disease, and associate with DENV-specific antibody responses, they do not appear to have a protective role.

## Introduction

Dengue viral infections represent one of the most rapidly emerging mosquito borne viral infections in the world, resulting in 390 million infections annually, with an estimated annual global cost of $8.9 billion [1] [2]. The WHO has named dengue as one of the ten threats to global health in 2019 [3]. Intense disease monitoring with meticulous fluid management is currently the only management option, as specific treatments options are not yet available. The only registered dengue vaccine, was shown to have poor efficacy against some DENV serotypes and has a potential to increase the risk of severe disease in dengue seronegative individuals [4–6]. Although infection with a particular serotype of the DENV is thought to result in lifelong immunity, there have been instances where infection with the same serotype occurs despite the presence of neutralizing antibodies specific for that serotype [7, 8]. In addition, some individuals who had high neutralizing antibody titres to DENV2 following vaccination, subsequently developed infection with the same serotype [9]. Therefore, as the correlates of protection against the DENV are still not clear, it is essential to understand the role of T cells and antibodies in the protection against dengue infection.

The role of both conventional CD4+ and CD8+ T cells in dengue has been well studied and show they are highly activated, [10–13] and recent investigations have shown that T cells in dengue are likely to have a protective role [10, 12]. However, apart from eliminating cells infected by the DENV and production of antiviral cytokines, certain subsets of T cells such as the follicular helper T cells (Tfhs) also determine the type and magnitude of development of virus specific antibody responses. Tfh cells, which are located in the germinal centre, play a crucial role in activating germinal centre B cells leading to activation, isotype switching and production of long lasting neutralizing antibodies [14, 15]. Some Tfh cells are also found in the peripheral circulation and are identified by the expression of CXCR5. They also upregulate PD-1, ICOS, Bcl-6, CD40 and produce IL-21, which are important for the activation of germinal centre B cells[15]. It was recently shown that Tfh cells are highly activated in acute dengue and also correlated with the frequency of plasmablasts [16]. The Tfh cells were also shown to be able to produce IL-21 and IFNγ upon stimulation with DENV peptides, suggesting that they were capable of providing B cell help, which is largely dependent on production of IL-21[17]. Therefore, increase in frequency and activation of Tfh cells could lead to higher DENV specific antibody levels which may lead to protection, or possibly disease enhancement [18, 19]. In order to further understand the pathogenesis of acute dengue and the constituents of a protective immune response, it would be important to study how the functionality and phenotype of Tfh cells associate with DENV-specific antibody responses and clinical disease severity.

Dendritic cells (DCs) are crucial in activating naïve CD4+ T cells and inducing their differentiation to different CD4+ subsets[20]. DENV was shown to activate both RIG-I and MDA5 resulting in production of IFNβ and STAT1 phosphorylation, which led to production of IL-27 by both monocyte-derived and skin-derived DCs [21]. This in turn was shown to lead to polarization of naïve CD4+ T cells into Tfh cells by DENV infected DCs [21]. The Tfh cells activated B cells, and indeed a massive plasmablast expansion has been seen in patients with acute dengue [22, 23]. Monocyte, which are important antigen presenting cells are one of the main cell types infected by the DENV during acute illness [24, 25]. Recently we showed that monocytes of healthy individuals with past severe dengue responded to infection with the DENV by producing higher viral loads, increased levels of inflammatory cytokines and altered gene expression compared to monocytes of healthy individuals who had past inapparent dengue infection [26]. Therefore, it is possible that monocytes and DCs of those who are likely to develop severe dengue, cause increased activation and differentiation of Tfc cells to produce protective or pathogenic virus specific antibodies.

In this study, we investigated the phenotype and functionality of Tfh cells in acute dengue in patients with varying severity of acute dengue and studied the association of Tfh responses with plasmablast expansion, DENV specific IgM and IgG antibodies, NS1 specific antibodies and DENV serotype specific neutralizing antibody titres to further understand how the functionality and phenotype of Tfh cells, associate with expansion of plasmablasts and DENV specific antibody responses in acute dengue infection.

## Results

Of the 40 patients included in the study to investigate the functionality and phenotype of Tfh cells, the mean duration of illness at the time of recruitment was 7.1 (SD ± 1.9) days. Their mean age was 35.98 years (SD ± 12.3). 18 patients had DF and 22 had DHF and all these patients had a similar duration of illness. One of the patients with DHF developed shock. Mild bleeding manifestations such as epistaxis and gum bleeds were seen in five patients. All the samples were obtained from patients with DHF and DF at the same time (mean day of illness 7.1) and therefore, samples were obtained from those with DHF while they were in the critical phase. All patients with DF and DHF were followed throughout the illness and none of the patients with DF later developed fluid leakage and progressed to develop DHF. The clinical and laboratory features of these patients is described in Supplementary table 1. All had an acute DENV2 infection. According to the results of the commercial capture-IgM and IgG ELISA, 13/40 had primary infection while the remaining 27 had secondary infection.

### Expansion of Tfhs during acute dengue illness

As CXCR5 is a key surface molecule expressed on Tfh cells, its expression has been used as an important marker of Tfhs [17]. Therefore, we identified Tfh cells as CXCR5 expressing CD4+ T cells for identification of peripheral Tfhs as described previously [17, 27]. As CXCR5 expression was shown to be altered upon stimulation of T cells [17], the Tfhs were identified *ex vivo* without any stimulation and the intracellular cytokine staining (ICS) for IL-21 was also carried out *ex vivo* without stimulation. We further phenotyped the Tfhs based on the expression of PD-1, ICOS and IL-21 (Supplementary fig. 1). Representative flow cytometry plots for total Tfhs and IL-21 producing Tfhs in a patient with acute DENV infection are shown in figures 1a and 1b, respectively.

**Figure 1:**
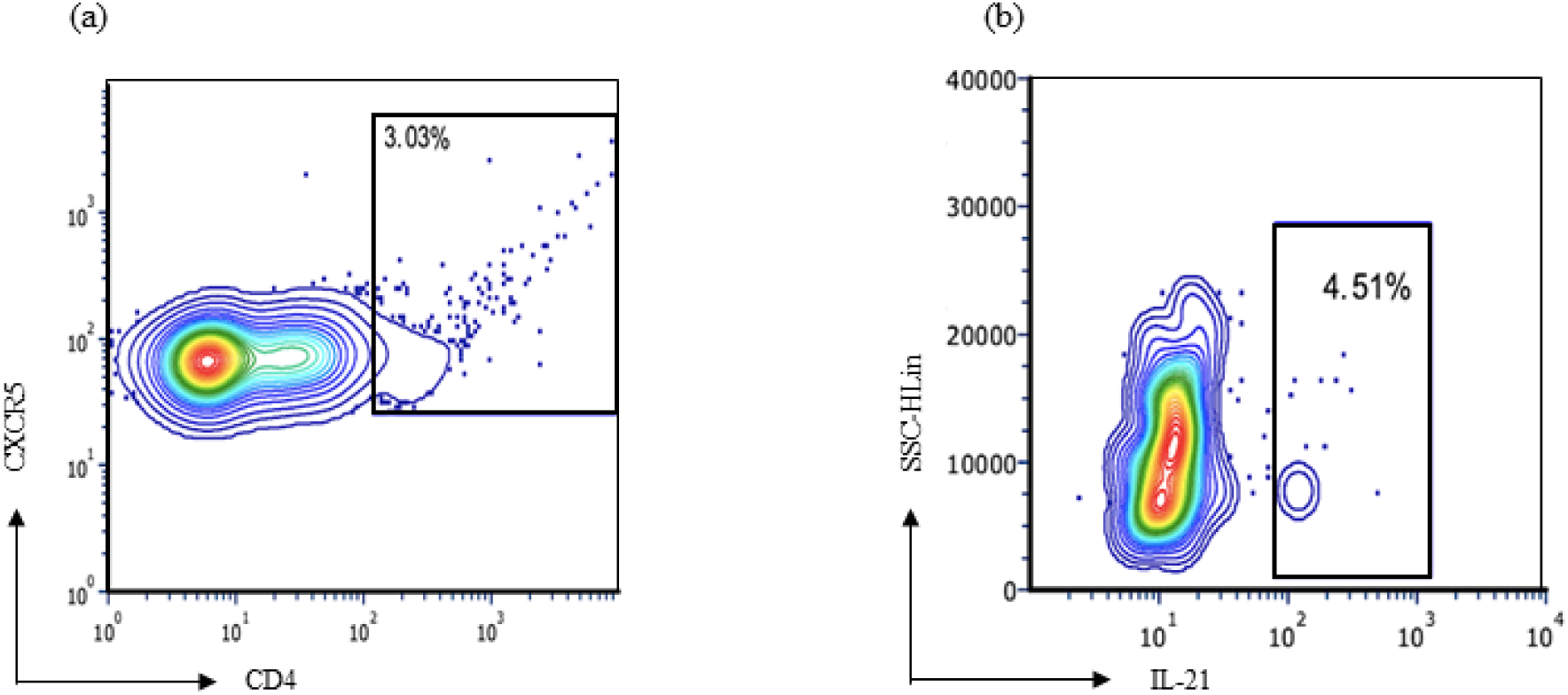
The frequency of Tfhs and IL-21 producing Tfhs in a patient with acute dengue in the acute phase. The expression of (a) total Tfhs (cells expressing CD4+CXCR5+) and (b) IL-21 producing Tfhs in a patient with acute dengue infection.

The frequency of Tfhs in patients with acute illness (median = 4.21%, IQR = 3.49 to 5.69 of CD4+ T cells) was significantly higher (p<0.0001) when compared to the frequencies in healthy individuals (median = 2.13%, IQR = 1.10 to 2.82 of CD4+ T cells). The frequency of Tfhs was also significantly higher (p<0.0001) in patients with DHF (median = 5.51%, IQR = 4.84 to 5.97 of CD4+ T cells) than in patients with DF (median = 3.59%, IQR = 2.99-3.95 of CD4+ T cells). In addition, a significant expansion (p = 0.001) was observed in patients with secondary dengue infection (median = 5.33%, IQR = 2.70 to 5.92 of CD4+ T cells) compared to those with primary dengue infection (median = 3.67%, IQR = 3.01 to 3.90 of CD4+ T cells) (Fig 2). 9 patients with DF had a primary infection, while 9 had a secondary dengue infection. There was no difference in the Tfh cell frequency in those with primary or secondary dengue infection, who had DF (p = 0.66). Furthermore, 4 patients with DHF had a primary infection, while 18 had a secondary dengue infection. In patients who had DHF, the frequency of Tfhs was significantly higher (p = 0.03) in those with secondary dengue infection (median = 5.64%, IQR = 5.28 to 6.06 of CD4+ T cells) compared to those with primary dengue infection (median = 3.40%, IQR = 3.08 to 5.35 of CD4+ T cells).

**Figure 2:**
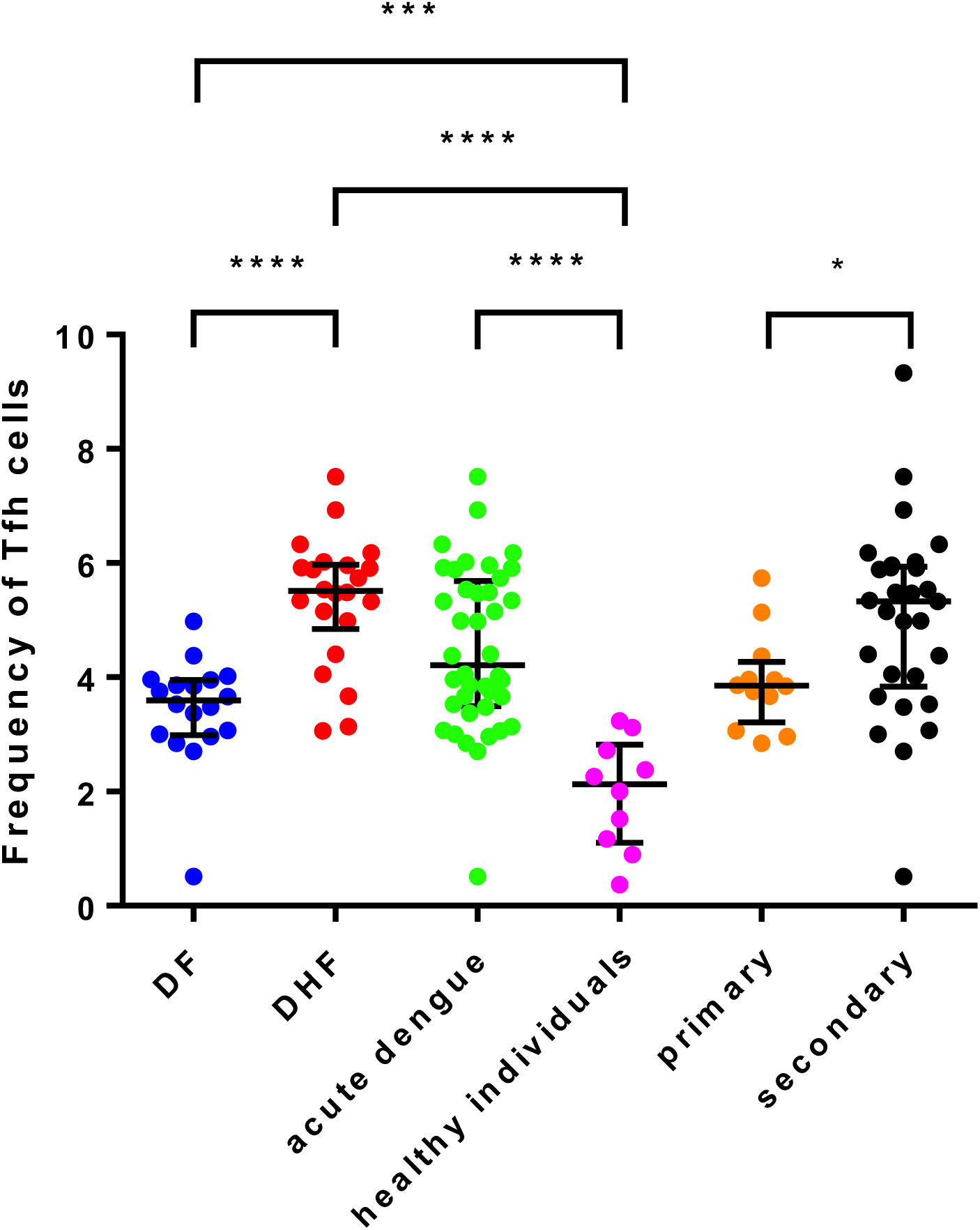
The frequency of Tfhs in patients with acute dengue. The frequency of Tfhs was assessed in patients with DF (n=18), in patients with DHF (n=22), in healthy individuals (n=12) in patients with primary infection (n=13) and secondary dengue infection (n=27) by multiparametric flowcytometry. The lines display the median and the inter quartile ranges. **P<0.01, ***p<0.001, ****p<0.0001

### Phenotypic analysis of Tfhs in acute dengue

PD-1 and ICOS have important functions in the development, functionality and migration of Tfhs to the germinal centres and are upregulated upon activation [15, 28]. Tfhs which express both PD-1 and ICOS are shown to be the most efficient helpers of the germinal centre B cells in antibody production [15]. In our study, PD-1 and ICOS co-expressing Tfhs were significantly expanded (p<0.0001) in patients with acute dengue (median = 8.28%, IQR = 2.96 to 12.59 of CD4+CXCR5+ T cells) compared to healthy individuals (median = 1.29%, IQR = 0.00 to 3.19 of CD4+CXCR5+ T cells). PD1+ICOS+ Tfhs were also significantly higher (p=0.0009) in patients with DHF (median = 11.03%, IQR = 8.11 to 12.72 of CD4+CXCR5+ T cells) compared to patient with DF (median = 2.81%, IQR = 2.17 to 7.27 CD4+CXCR5+ T cells) (Fig 3a).

**Figure 3:**
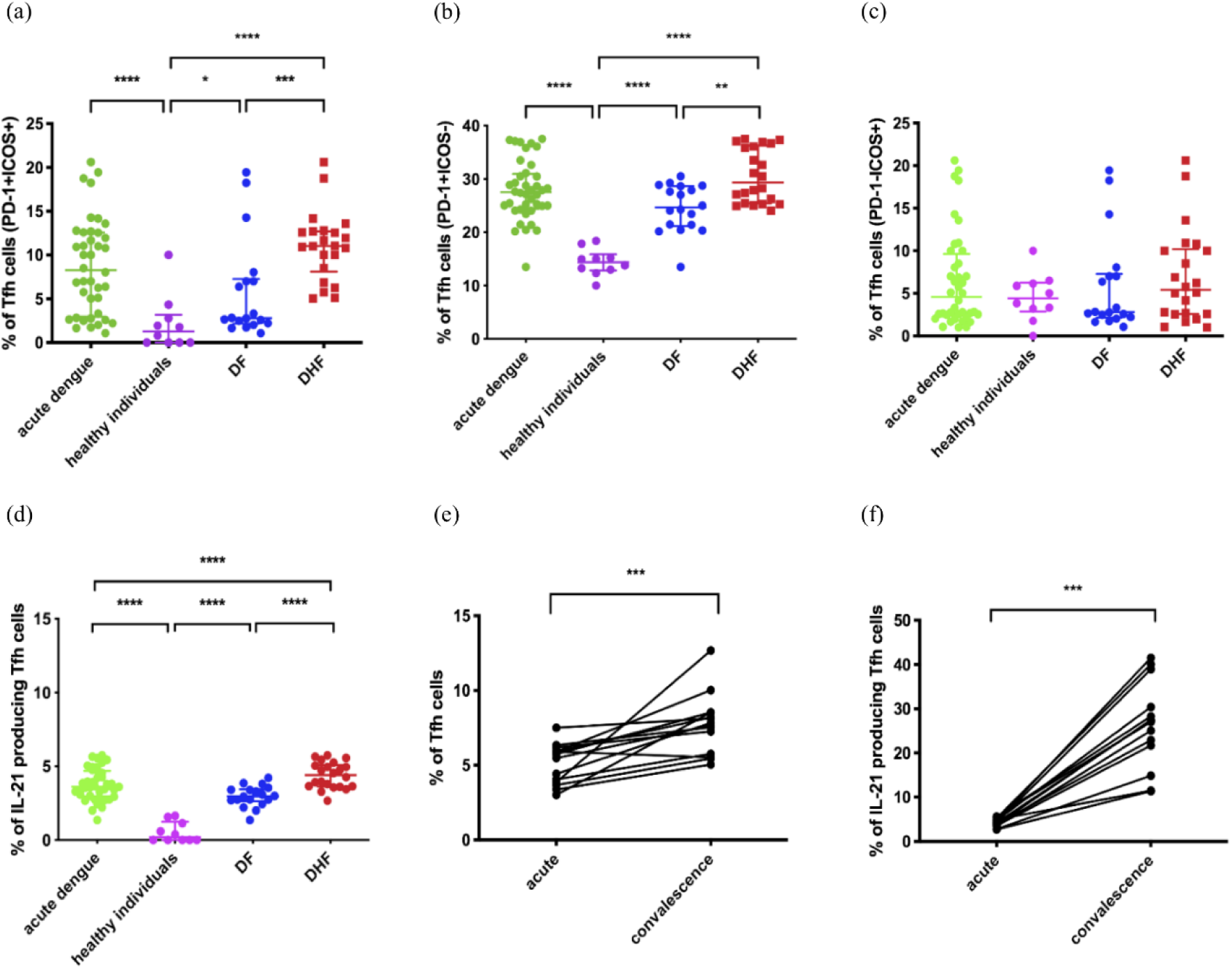
Phenotypical and functional differences of Tfhs in patients with acute dengue. The frequency of PD-1+ICOS+ co-expressing Tfhs (a) the frequency of Tfhs expressing PD-1+ alone (b) the frequency of Tfhs expressing ICOS alone (c) the frequency of IL-21 producing Tfhs in patients (d) is shown. The frequency of the total Tfhs and (e) and IL-21 producing Tfhs (f) were also assessed in a subset of these patients (n=14) during the acute and the convalescent phase of illness. The lines display the median and the inter quartile ranges. *P<0.05, **P<0.01, ***p<0.001, ****p<0.0001

The majority of Tfhs in patients with acute dengue only expressed PD-1 and not ICOS. Again, the PD-1+ICOS-Tfhs were significantly increased (p<0.0001) in patients with acute dengue infection (median = 27.51%, IQR = 24.45 to 30.95 of CD4+CXCR5+ T cells) compared to healthy individuals (median = 14.39%, IQR = 12.84 to 15.88 of CD4+CXCR5+ T cells) (Fig 3b). The frequency of PD-1+ICOS-Tfhs was also significantly higher (p=0.001) in patients with DHF (median = 29.36%, IQR = 25.41-36.24 of CD4+CXCR5+ T cells) compared to those with DF (median = 24.65%, IQR = 21.19 to 28.68 of CD4+CXCR5+ T cells) (Fig 3b). In contrast, the frequency of PD-1-ICOS+ Tfhs were not significantly different in acute infection (p = 0.70) compared to healthy individuals and in patients with DHF and DF (p=0.43) (Fig 3c).

### Functionality of Tfhs during acute dengue

Tfhs produce high levels of IL-21, a cytokine that is critical for germinal center formation and for inducing isotype switching of antibodies produced by germinal centre B cells[14]. Therefore, in order to investigate the cytokine production potential of Tfhs in acute dengue, we sought to determine IL-21 production by the Tfhs in patients with acute dengue *ex vivo*. The frequency of IL-21 producing Tfhs were highly significant (p<0.0001) in patients with acute dengue (median = 3.62%, IQR = 2.95 to 4.67 CD4+CXCR5+ T cells) when compared with healthy individuals (median = 0.19%, IQR = 0.00 to 1.24 CD4+CXCR5+ T cells), where hardly any cytokine production was observed (Fig 3d). The frequency of IL-21 producing Tfhs were also significantly higher (p<0.0001) in patients with DHF (median = 4.41%, IQR = 3.63 to 5.10 CD4+CXCR5+ T cells) compared to those with DF (median = 2.96%, IQR = 2.63 to 3.43 CD4+CXCR5+ T cells) (Fig 3d).

As the frequency of Tfhs were assessed in the acute phase, we proceed to investigate the phenotype and functionality of these cells in the convalescent period (21 to 30 days since onset of illness) in 14/40 in this cohort. Interestingly, the frequency of Tfhs was significantly higher (p = 0.0002) during the convalescent phase in these patients (median = 7.74%, IQR = 5.70 to 8.53 of CD4+ T cells) compared to the frequencies in acute infection (median = 5.60%, IQR = 3.81-5.98) (Fig. 3e). Furthermore, the frequency of IL-21 producing Tfhs was significantly higher (p = 0.0001) during convalescent phase (median = 27.15%, IQR = 19.97 to 32.52 CXCR5+CD4+ cells) compared to acute phase of illness (median = 4.24%, IQR = 3.52 to 5.02 CXCR5+CD4+ cells) (Fig 3f). However, the proportion of Tfh cells positive for PD-1 was significantly less (p=0.0004) in the convalescent phase (median 13.7 IQR 10.3% to 16.3%) compared to the acute phase (median 27.2, IQR 24.7% to 31.8%). Representative flow cytometry plots for total Tfhs and IL-21 producing Tfhs during the convalescent phase are shown in figures 4a and 4b, respectively.

**Figure 4:**
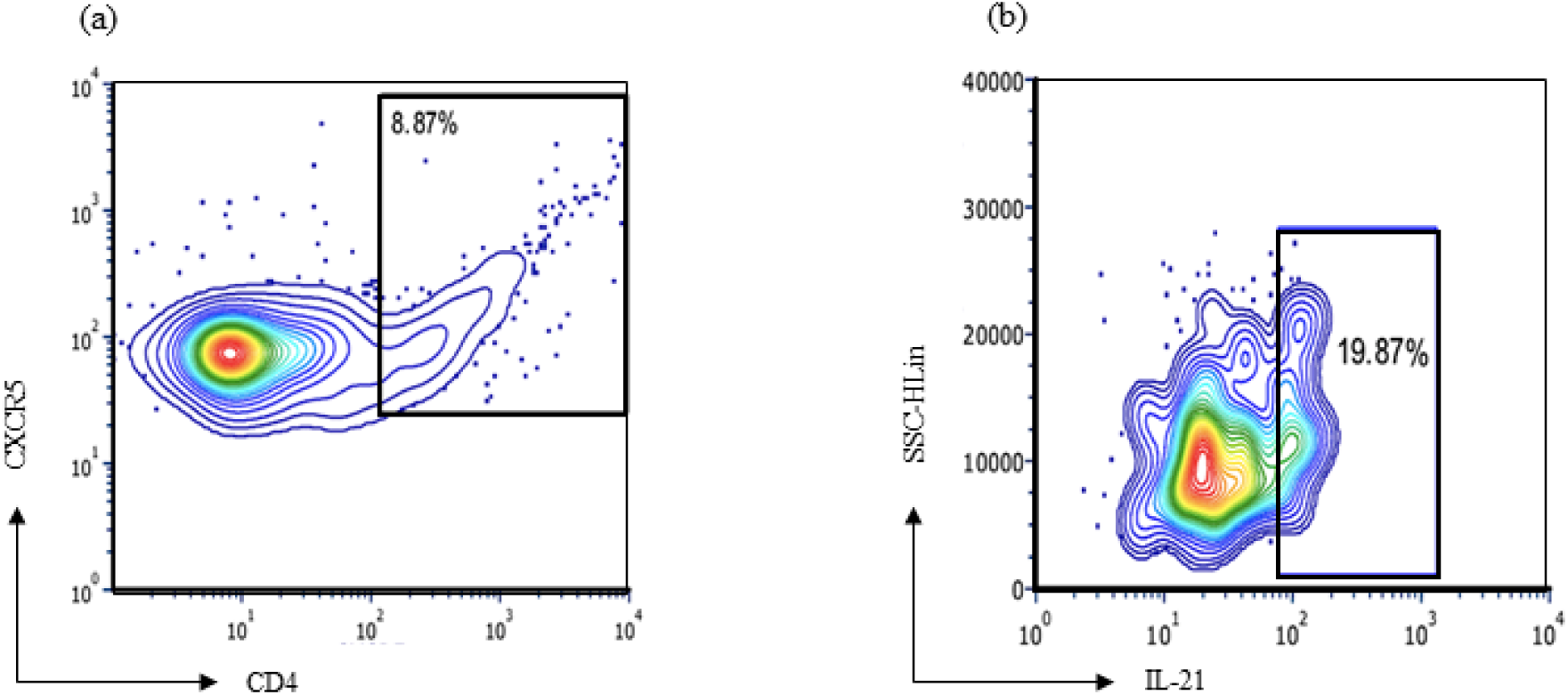
The frequency of Tfhs and IL-21 producing Tfhs in a patient with acute dengue in the convalescent phase. The expression of (a) total Tfhs(cells expressing CD4+CXCR5+) and (b) IL-21 producing Tfh cells in a patient during the convalescent phase (same patient as shown in Figure 1).

### Association of Tfhs with virological and laboratory parameters

The frequency of Tfhs inversely correlated with the viral loads (Spearman r = −0.63, p<0.0001) of patients with acute dengue (Table 1), suggesting that Tfh expansion was associated with resolution of viraemia. Although a significant inverse correlation was seen with the frequency of IL-21 producing Tfhs and the viral loads (Spearman’s r = −0.57, p = 0.0002) (Table1), no correlation was observed between PD1^+^ICOS^+^, PD1^+^ICOS^−^, PD1^−^ICOS^+^ subsets of Tfh cells. The frequency of Tfhs showed an inverse correlation with platelet counts in patients with acute dengue infection (Spearman r = −0.42, p = 0.008). However, the platelet counts did not correlate with the frequency of IL-21 producing Tfhs (Spearman r = −0.31, p = 0.058).

**Table 1:**
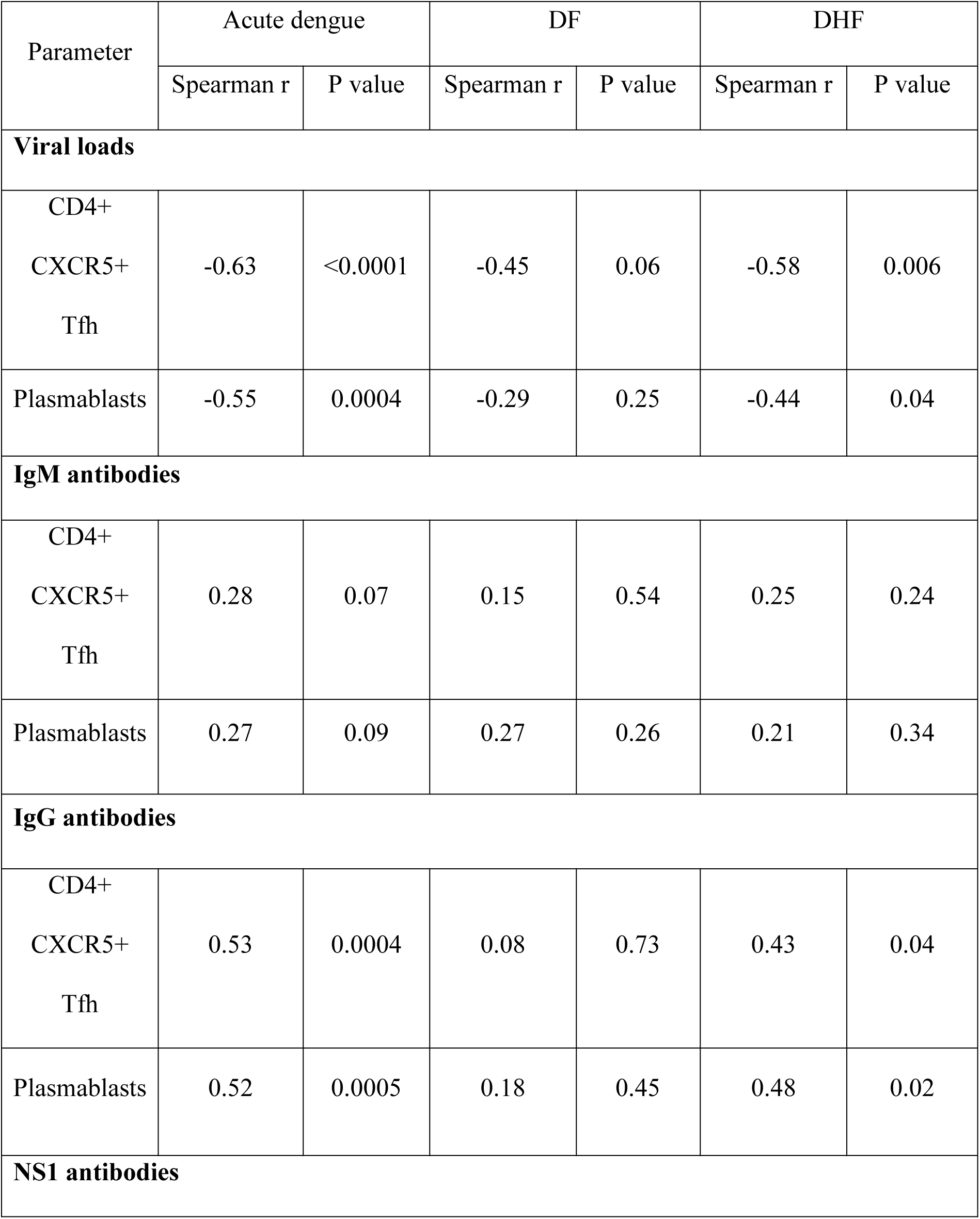
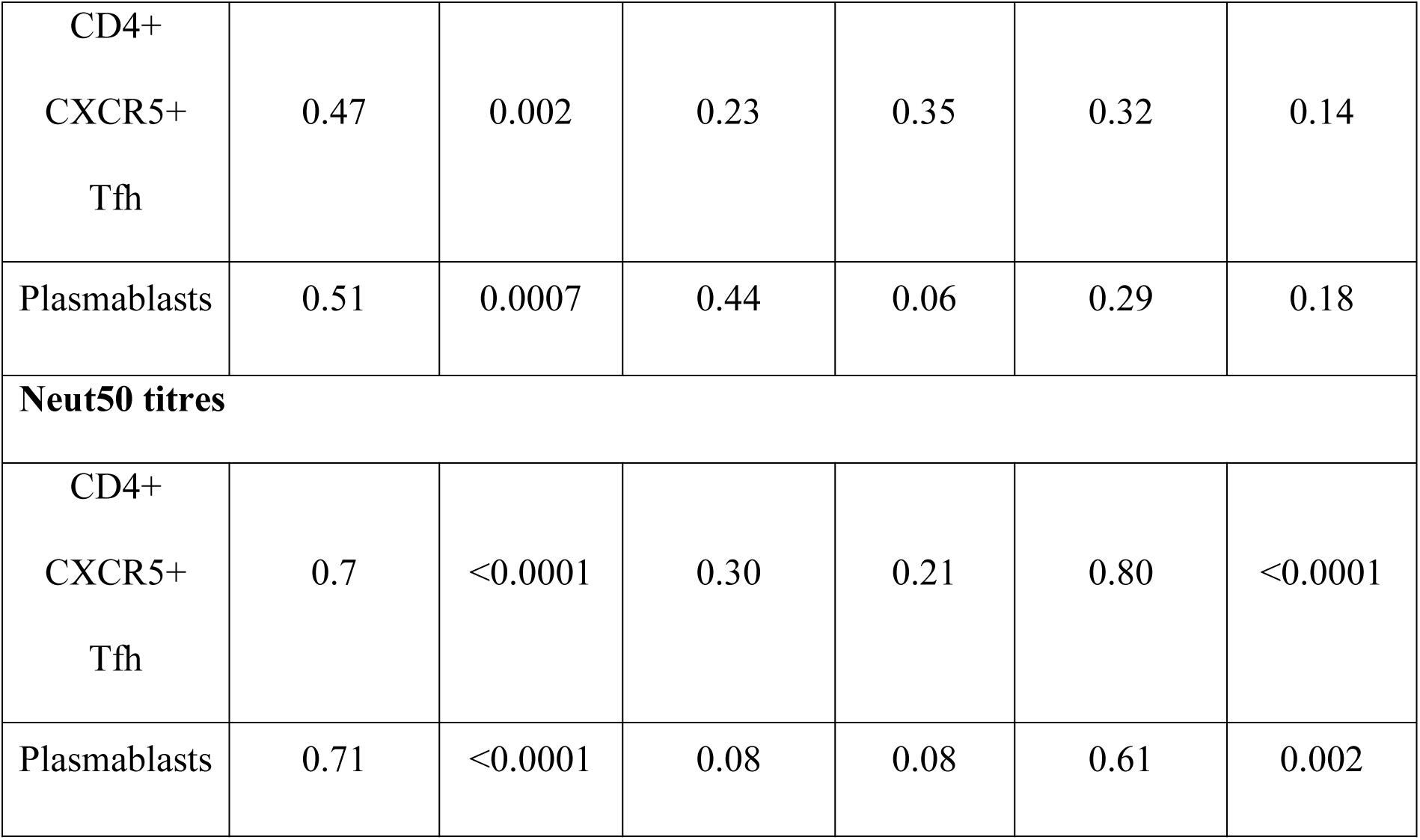
Association of Tfhs and plasmablasts with viral loads and DENV specific antibody responses

### Plasmablast expansion in acute dengue infection and correlation of frequency of plasmablasts with frequency of Tfhs

A massive expansion of plasmablasts has been observed in acute dengue infection and the activated Tfhs have been shown to correlate with the frequency of plasmablasts [16, 22]. Since Tfhs activate germinal centre B cells and Tfh cytokines such as IL-21 induce proliferation of plasmablasts, we investigated the association of the Tfh activation and cytokine production with the frequency of plasmablasts (Supplementary fig. 2). A representative flow cytometry plot for plasmablasts in a patient with acute infection and during convalescence is shown in figure 5. As shown previously [16, 22], we too found that plasmablasts significantly expanded (p<0.0001) during acute infection (median = 30.32, IQR = 26.24 to 42.73 of total B cells) compared to healthy individuals (median = 0.87, IQR = 0.40 to 1.50 of total B cells) (Fig 6a). The frequencies of plasmablasts associated with clinical disease severity as they were significantly higher (p<0.0001) in patients with DHF (median = 39.23, IQR = 30.47 to 48.26 of total B cells) when compared to those with DF (median = 26.47, IQR = 20.54 to 29.72 of total of B cells) (Fig 6a). The frequency of plasmablasts was also significantly higher (p=0.03) in patients with secondary dengue infections (median = 34.51, IQR = 29.87 to 44.83 of B the total B cells) to those with primary infections (median = 26.98%, IQR = 22.19 to 35.70 of the total B cells) (Fig 6a).

**Figure 5:**
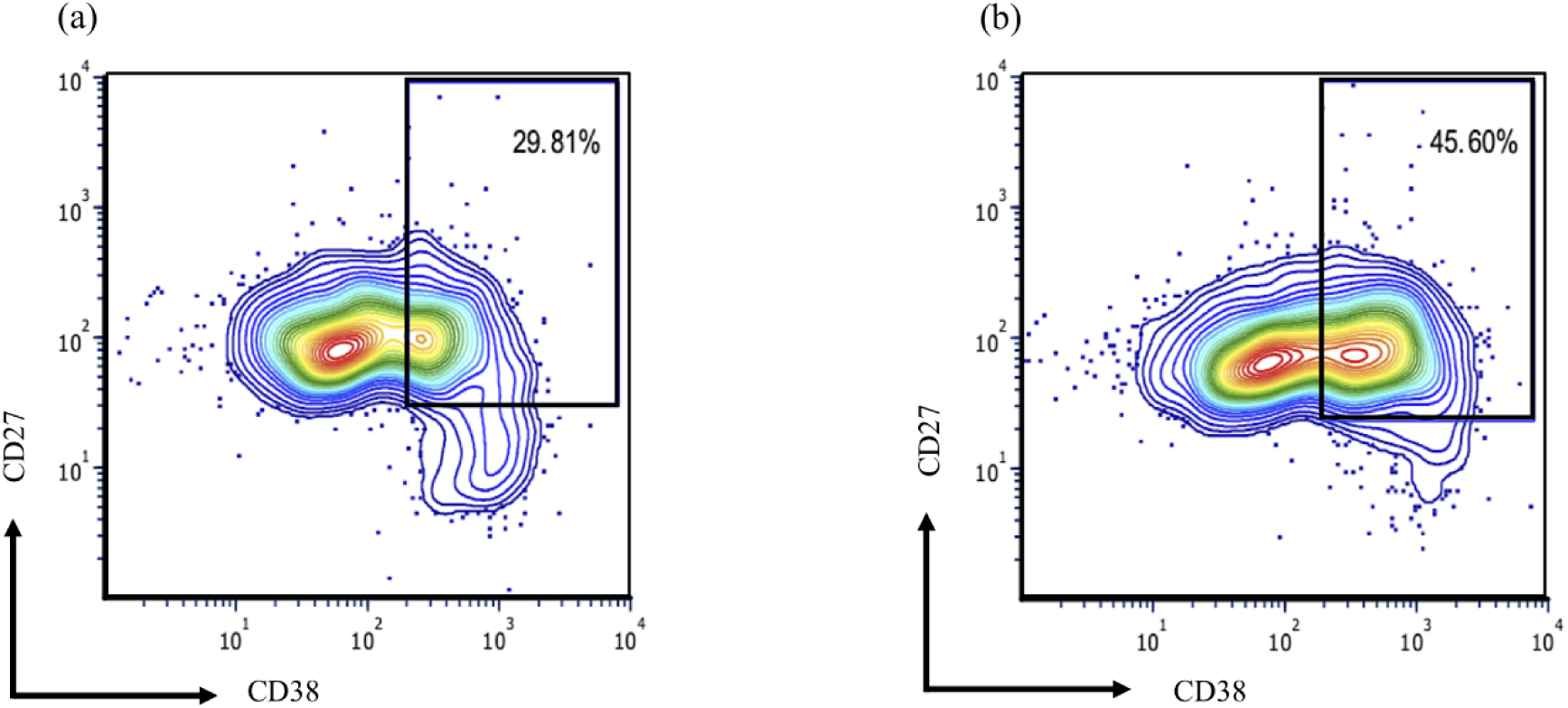
The frequency of plasmablasts in a patient with acute dengue in the acute and convalescent phase. The expression of plasmablasts (CD19 cells expressing CD38 and CD27) are shown in the same patient during the acute (a) and convalescent phase (b).

**Figure 6:**
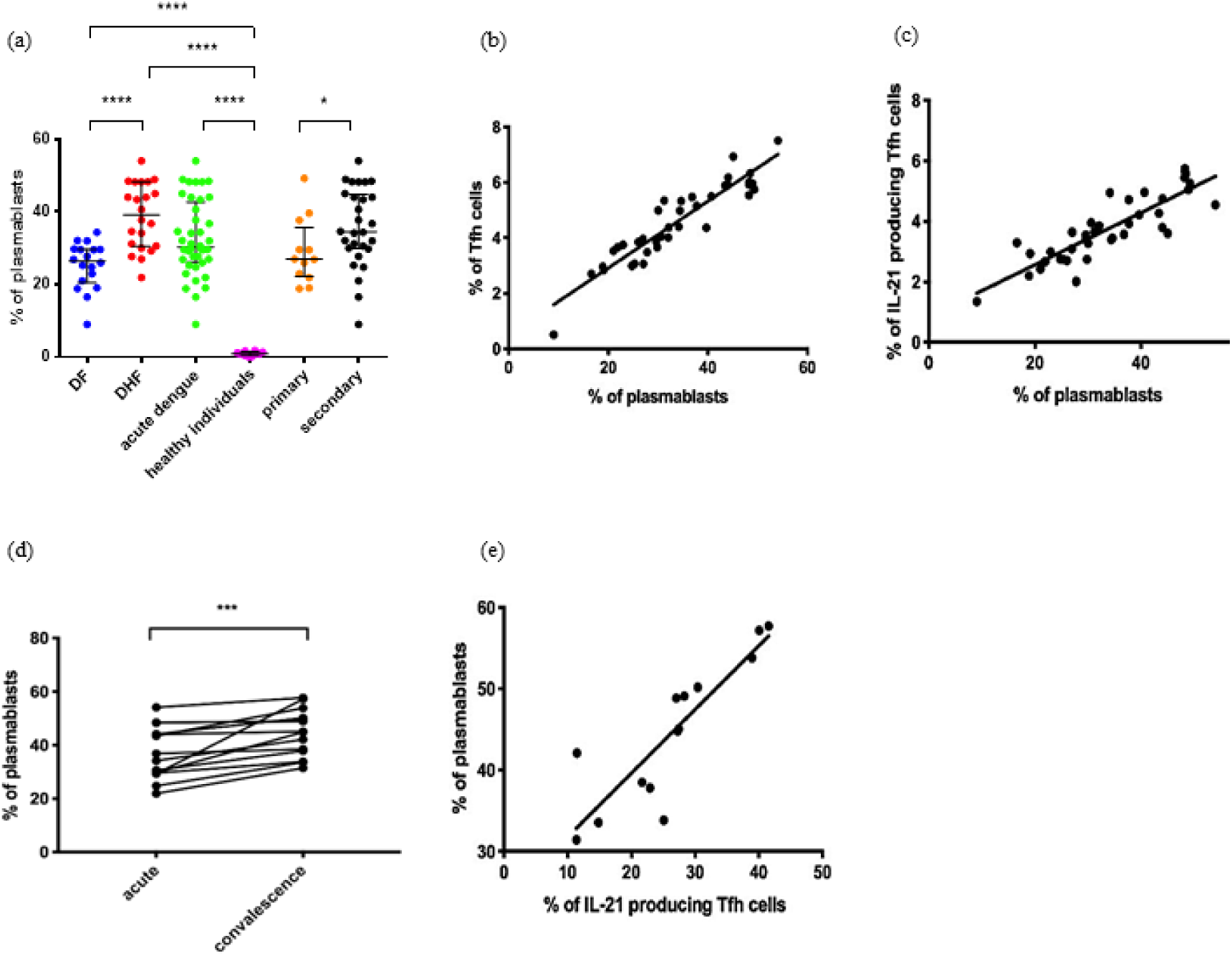
Plasmablast expansion in acute dengue infection and correlation of frequency of plasmablasts with frequency of Tfhs. (a) frequency of plasmablasts in patients with DF and DHF, primary and secondary infections, acute dengue and healthy individuals (b) correlation of frequency of plasmablasts with frequency of Tfhs in acute infection (Spearman r = 0.91) (c) correlation of frequency of plasmablasts with frequency of IL-21 producing Tfhs in acute infection (Spearman r = 0.78) (d) frequency of plasmablasts during the acute phase of the illness and during the convalescent phase. (e) correlation of frequency of IL-21 producing Tfhs with frequency of plasmablasts during the convalescent phase (Spearman r = 0.92). The lines display the median and the inter quartile ranges. *P<0.05, ***p<0.001, ****p<0.0001

The frequency of plasmablasts significantly and positively correlated with the frequency of the total Tfh cells (Spearman R=0.91, p<0.0001) (Fig 6b). Although the frequency of plasmablasts correlated with Tfh cells expressing PD1+ICOS+ (Spearman r=0.37, p=0.02) and with PD1+ICOS-(Spearman r = 0.34, p=0.03) the greatest correlation was observed with Tfh cells producing IL-21 (Spearman r = 0.78, p<0.0001) (Fig 6c). A significant inverse correlation was seen between the frequency of plasmablasts and the viral loads (Spearman r = −0.55, p = 0.0004) as well as with the lowest platelet count in the patients (Spearman r = −0.38, p = 0.01) (Table 1).

As with the frequency of Tfhs, we also determined the frequency of plasmablasts in the convalescent phase in 14/40 patients in this cohort. The frequency of plasmablasts was significantly higher (p = 0.0001) in convalescent phase (median = 44.95%, IQR = 36.84 to 51.10 of total B cells) than in the acute infection (median = 35.48%, IQR = 29.50 to 45.13 of total B cells) (Fig. 5 and 6d). Similar to the observations in the acute phase of illness, the frequency of plasmablasts significantly correlated with the frequency of IL-21 secreting Tfhs in the convalescent phase (Spearman r = 0.92, p<0.0001) (Fig. 6e).

### Association of Tfhs and plasmablasts with DENV envelope protein specific antibody responses

The majority of the plasmablast response was found to be specific for the DENV in acute dengue, predominantly comprising IgG secreting cells [22]. However, in a previous study no association was seen between the frequency of plasmablasts and DENV-specific neutralizing antibody titres (Neut50) [23]. Therefore, in order to study the relationship between Tfhs, plasmablasts and DENV specific antibodies we investigated the association of Neut50, total DENV antibodies and also DENV NS1 specific antibodies in this cohort of patients.

We semi-quantitatively measured DENV-specific IgM and IgG antibody levels in serum (expressed as Panbio units), using the PanBio IgM and IgG capture ELISA, which uses the DENV envelope protein as the coating antigen. There was no significant difference (p = 0.42) in DENV-specific IgM titres between patients with DF (median = 26.23 Panbio units, IQR = 9.22 to 40.62) and DHF (median = 33.83 Panbio units, IQR = 18.69 to 42.66), whereas DENV-specific IgG titres were significantly higher (p = 0.004) in DHF patients (median = 47.17 Panbio units, IQR = 41.03 to 50.09) than in those with DF (median = 8.17 Panbio units, IQR = 0.79 to 29.62). We detected no significant correlation between DENV-specific IgM antibody titres and frequency of Tfhs in patients with DF (Spearman r = 0.15, p = 0.54) and DHF (Spearman r = 0.25, p = 0.24) (Table 1). As observed with Tfh cells, DENV-specific IgM antibody titres did not correlate with the frequency of plasmablasts in patients with DF (Spearman r = 0.27, p = 0.26) or in DHF (Spearman r = 0.21, p = 0.34) (Table 1).

The DENV-specific IgG titres also did not correlate with the frequency of Tfhs in patients with DF (Spearman’s r = 0.08, p = 0.73), but a significant correlation was seen in patients with DHF (Spearman’s r = 0.43, p = 0.04) (Table 1). Similar findings were observed with plasmablasts with no significant correlation between DENV-specific IgG titres in DF patients (Spearman r = 0.18, P = 0.45) and significant correlation in patients with DHF (Spearman r = 0.48, p = 0.02) (Table 1). Furthermore, a significant correlation was detected between DENV-specific IgG antibody titres and frequency of IL-21 producing Tfhs in patients with acute dengue (Spearman r = 0.39, p = 0.01).

### Association of Tfhs and plasmablasts with DENV NS1-specific antibody responses

Although the majority of DENV specific antibodies are specific to the DENV envelope protein, 27% of the DENV antibody repertoire has shown to be specific to DENV NS1 protein [29]. Therefore, we measured DENV2 specific NS1 antibodies using an in-house ELISA as described before [18]. As observed earlier, NS1-antibody levels were significantly higher in patients with DHF (median 1.7, IQR 0.76 to 2.3 OD values) compared to those with DF (median 0.56, IQR 0.23 to 1.4 OD values). We observed a significant correlation between NS1 antibody levels and frequencies of only IL-21 producing Tfh cells in DF (Spearman r = 0.48, p = 0.04) but not in patients with DHF (Spearman r = 0.05, p = 0.81). However, a significant correlation was not observed between NS1 antibody levels and frequencies of Tfh cells in patients with DF (Spearman r = 0.23, p = 0.35) and DHF (Spearman r = 0.32, p = 0.14). Similarly, we did not observe a significant correlation between NS1 antibody levels and frequencies of plasmablasts in both DF (Spearman r = 0.44, p = 0.06) and in DHF (Spearman r = 0.29, p = 0.18) (Table 1). As shown in our earlier cohorts, the NS1 antibody levels inversely correlated with platelet counts (Spearman r=-0.38, p=0.02) and with viral loads (Spearman r=-0.5, p=0.001).

### Association of Tfhs and plasmablasts with DENV specific neutralizing antibody titres

A significant inverse correlation was seen in the viral loads with DENV-specific IgG titres (Spearman r = −0.34, P = 0.03) and with DENV-specific NS1 antibody levels (Spearman r = - 0.48, p = 0.002). However, as the Panbio ELISA measured all DENV envelope specific antibodies and not necessarily neutralizing antibodies, which are thought in some studies to associate with protection, we then proceeded to determine association of neutralizing antibodies specific to the infecting DENV serotype (DENV2) with the frequency of Tfhs and plasmablasts. The Neut50 titres were significantly higher (p = 0.006) in patients with DHF (median = 35821, IQR = 16,393 to 91,086) than in the patients with DF (median = 5314, IQR = 2586 to 12,343). Although there was no significant correlation in Neut50 titres with the frequency of Tfhs in patients with DF (Spearman r = 0.30, p = 0.21), a significant correlation was detected between Neut50 titres and the frequency of Tfhs in patients with DHF (Spearman’s r = 0.80, p<0.0001) (Table 1). We observed the same relationship with the frequency of plasmablasts. i.e. there was no significant correlation between Neut50 titres and the frequency of plasmablasts in patients with DF (Spearman r = 0.42, p = 0.08) but in patients with DHF (Spearman r = 0.61, p = 0.002) (Table 1). Interestingly, in this cohort of patients who all had a DENV2 infection, Neut50 titres only inversely correlated with the viral loads in patients with DF (Spearman = −0.75, p = 0.0003) and not in those with DHF (Spearman’s r = - 0.26, p=0.23) (Fig 7a and 7b).

**Figure 7:**
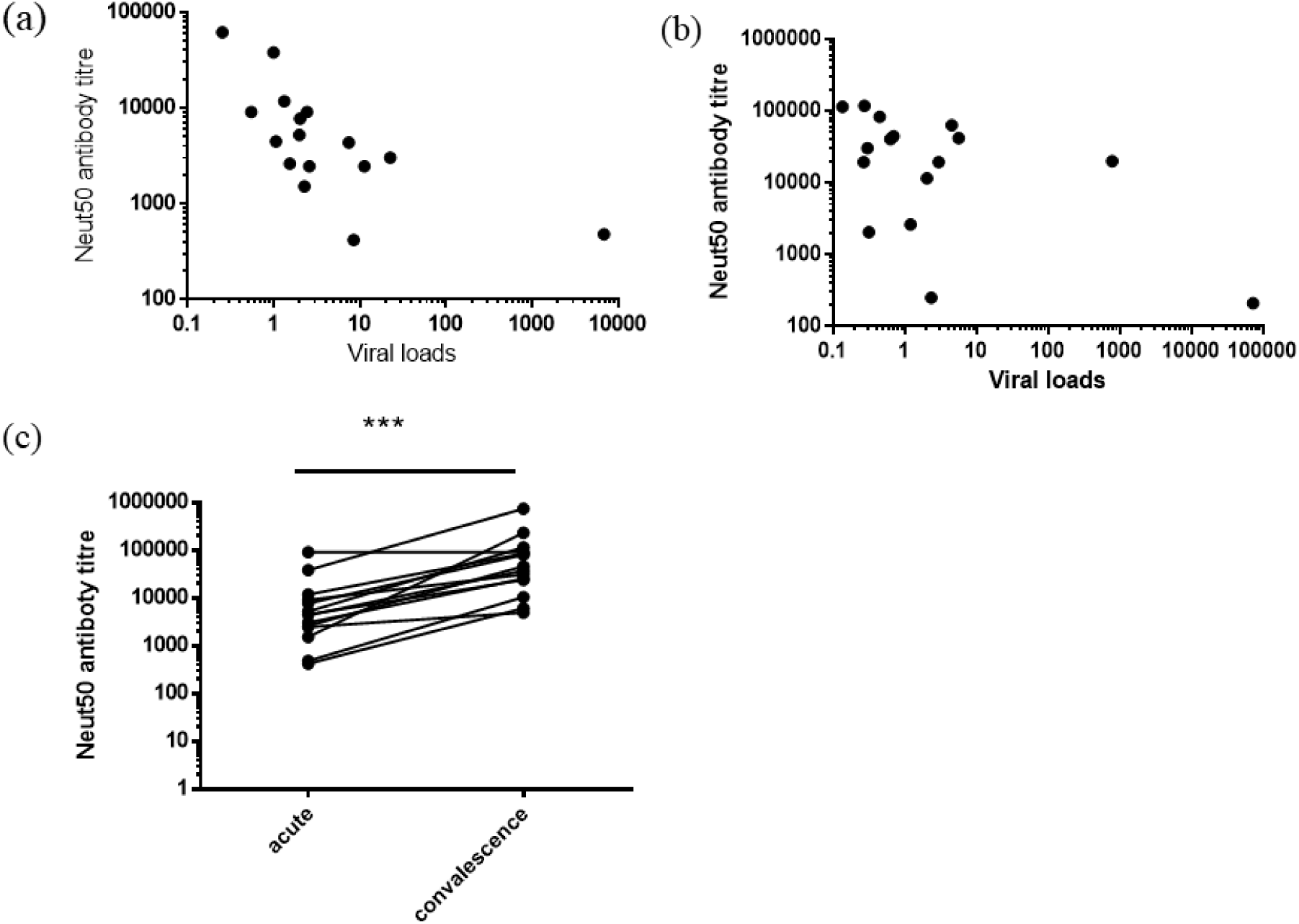
Association of neutralizing antibody titres (Neut50) with viraemia and changes with time. The Neut50 titres were measured in patients with DF (n=18) and DHF (n=22) during the acute phase and correlated with the viral loads. (a) A significant and negative correlation of viral loads with Neut50 antibody titres were seen in patients with DF (Spearman r = −0.75, p=0.003) but not with DHF (Spearman r = −0.26, p=0.23) (b) Neut50 titres were also measured in patients during the acute phase and the convalescent phase in patients (n=18) (c). ***p<0.001

The Neut50 titres were measured in the convalescent phase (day 21 to 30 since onset of illness) in 14/40 patients with acute dengue, and the Neut50 titres were 10 fold higher (p = 0.0002) in the convalescent phase (median 41,432, IQR=20,609 to 97,105 Neut50 titres) when compared to the acute phase (median 4416, IQR 2,220 to 9,777 Neut50 titres) (Fig 7c). The Neut50 antibody titres showed a significant correlation with the IL-21 producing Tfh cells (Spearman r=0.73, p=0.004) and plasmablasts (Spearman r = 0.85, p=0.0002) but not with the total Tfh cells (Spearman r = 0.06, p=0.82).

### Kinetics of neutralizing antibody production in patients with varying severity of acute DENV1 and DENV2 infection

The above data show that Neut50 antibody titres significantly correlated with the frequency of Tfh cells and plasmablasts in patients with DHF but not in those with DF and also inversely correlated with the viral loads only in patients with DF. Although a Neut50 antibody titre of 1:10 is considered protective in some studies [30, 31], some individuals high neutralizing antibodies for a particular DENV serotype later went on to develop DHF when infected with that particular serotype[32]. Furthermore, those who were found to have high Neut50 titres to DENV2 in phase 2b and 3 dengue vaccine trials, were later found to be infected with the same serotype when naturally exposed to the virus [33, 34]. Since the association of Neut50 titres with clinical disease severity has not been assessed, we sought to investigate the changes in the Neut50 titres throughout the course of illness in patients with an acute DENV1 or an acute DENV2 infection.

As the Neut50 titres are likely to be different in those with primary and secondary dengue longitudinally, in order to compare disease severity, we investigated the kinetics of Neut50 titres in a cohort of patients with secondary dengue (this is a different cohort that the patients recruited for the Tfh studies). We studied the kinetics of the Neut50 titres in 40 patients with varying severity of acute DENV-1 infection (DF = 12, DHF = 10) and DENV-2 infection (DF = 8, DHF = 9) throughout the course of illness. Tthe first sample was obtained before any of the patients proceeded to develop either DHF or DF.

We found that the Neut50 titres of those with acute secondary DF and DHF in both DENV1 and DENV2 infection were similar in the febrile phase before the onset of the plasma leakage in patients with DHF (Fig 8a and 8b). While the Neut50 titres exponentially increased in patients with DF in acute secondary DENV1 infection, the opposite was observed in acute secondary DENV2 infection. However, there was no significant difference in the viral loads in patients with DF and DHF at any time point in the illness in acute secondary DENV1 infection (Fig 8c) or in acute secondary DENV2 infection (Fig 8d). The viral loads at the beginning of the illness did not correlate with the Neut50 titres seen in the course of illness in patients with DF or DHF in DENV1 infection at any timepoint measured. No such associations were also observed for either DF or DHF in acute secondary DENV2 infection.

**Figure 8:**
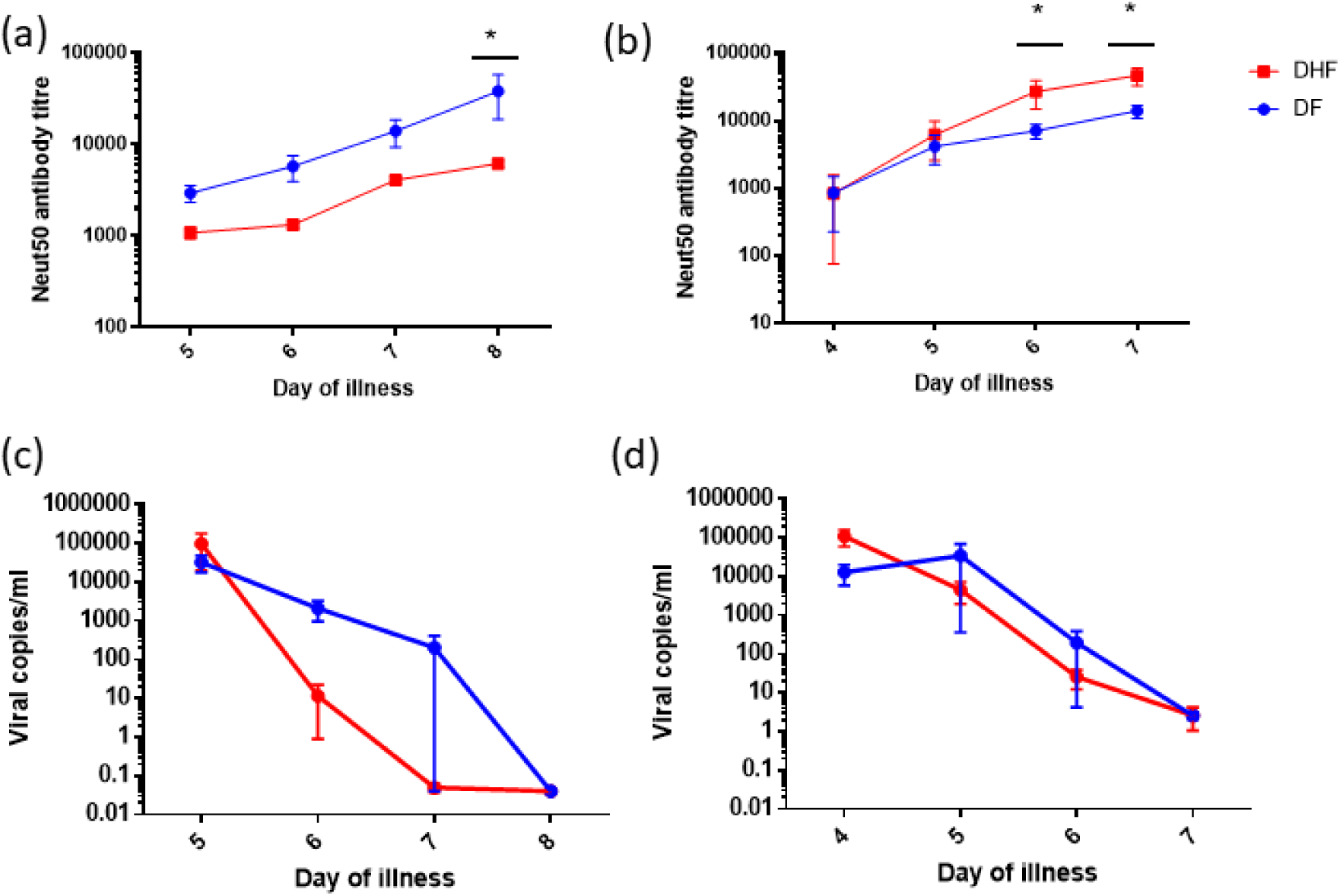
Kinetics and association of neutralizing antibody titres with viral loads. Neutralising antibody (Neut50) titres were assessed in acute secondary DENV1 infection in patients with DF (n=12) and DHF (n=10) (a) and in acute secondary DENV2 infection in patients with DF (n=8) and DHF (n=9) (b) throughout the course of illness from date of admission to discharge. Viral loads were also measured in the same cohort of patients with acute secondary DENV1 infection in patients with DF (n=12) and DHF (n=10) (c) and in acute secondary DENV2 infection in patients with DF (n=8) and DHF (n=9) (d) throughout the course of illness from date of admission to discharge.

At the end of the critical phase (in the recovery phase) the Neut50 antibody titres were several folds higher in those with both DF and DHF when infected with DENV2 compared to DENV1. For instance, those with an acute secondary DENV2 causing DHF had a median Neut50 titre of 43,652 (range 29,512 to 69,183 titre) compared to those with DF (median 14,234, range 8,913 to 19,055 titre). In contrast, those with an acute secondary DENV1 causing DHF had a median Neut50 titre of 5,890 (range 4,266 to 8,913) compared to those with DF (median 17,819, range 7,943 to 251, 188).

## Discussion

Our results show that Tfh cells are significantly expanded in acute dengue, especially in those with severe clinical disease (DHF). There was also a significant increase in the IL-21 producing Tfh cells in patients with DHF, which are the most functional subset of Tfh cells and therefore most capable of providing help to B cells [14]. The Tfhs co-expressing PD-1+ICOS+, which are the subtype of Tfh cells that are most efficient in inducing germinal centre B cell antibody production [15], were also significantly expanded, especially in those with DHF. Although all types of Tfh cells (PD-1+ICOS+, PD-1+ICOS-, PD1-ICOS+) correlated with the frequency of the plasmablasts, the IL-21 producing Tfh cells showed the most significant and the strongest correlation, suggesting that the most functional subset of Tfh cells was driving the plasmablast expansion in acute dengue infection. Although the Tfhs cells in general and the IL-21 producing Tfh cells further expanded in the convalescent phase along with plasmablasts, PD-1 expressing Tfhs cells significantly decreased suggesting that although the Tfh cells were functional, they were less activated during the convalescent phase. Production of neutralising antibodies too appear to continue after the acute phase (day 6 to 8 measured in this study) as the Neut50 titres were 10 times higher in the convalescent phase compared to the values in the acute phase. The Neut50 titres in the convalescent phase only correlated with the IL-21 producing Tfh cells, suggesting that this functional subset of Tfh cells is likely to be important to drive high titres of long-lasting neutralising antibody titres.

The total population of Tfh cells and plasmablasts significantly correlated with Neut50 antibody titres and DENV-specific IgG antibodies, indicating their importance in driving pathogen specific antibody responses during acute infection. Therefore, DENV specific-IgG, Neut50 titres and NS1-antibody levels were significantly higher in those with DHF compared to those with DF, probably due to increased numbers of plasmablasts producing higher levels of DENV-specific antibodies. The frequency of Tfh cells, plasmablasts, DENV specific IgG and Neut50 titres all inversely correlated with the viral loads, which suggest that they are likely to be important in the control of the virus. The Neut50 antibody titres were significantly higher in patients with DHF in the first cohort of patients infected with DENV2, which could be suggesting that increased viral loads in patients with DHF lead to increased expansion of Tfh cells and thus plasmablasts, resulting in higher Neut50 titres. However, we did not observe higher viral loads in patients with DHF compared to those with DF, in this cohort of 40 patients, whom we studied the Tfh, plasmablast and antibody responses. Since blood samples to study these responses were obtained on a mean duration of illness of day 7.1, we would have missed the early events that led to rise in the Neut50 titres in those with DHF compared to those with DF. In addition, although we did not observe differences in viral loads in patients with DHF compared to DF on day 6-8 of illness, it is possible that patients with high viral loads early in illness, subsequently progressed to develop DHF and also these higher early viral loads resulted in higher Neut50 titres in patients with DHF at a later time point in illness.

In order to address this question, we studied the kinetics of viral loads, Neut50 assays during the febrile phase in a different cohort of patients with acute secondary infection due to DENV1 and DENV2. We chose patients with a secondary dengue infection, as it would be unsuitable to compare antibody and viral loads in a cohort of patients with both primary and secondary dengue infection. We also chose patient cohorts infected with 2 different types of virus serotypes (DENV1 and DENV2) to better understand the changes in antibody and viral loads based on DENV serotype. Our data showed that there was no difference in the viral loads in patients with acute secondary DENV1 or DENV2 infection during the febrile phase, before any of the patients progressed to develop vascular leak and thus DHF. In addition, there was no difference in the viral loads throughout the illness in patients with DHF or DF due to infection with either viral serotype. However, in both DENV1 and DENV2 infections, while the viral loads continued to decline during the illness, the Neut50 titres rose.

During the febrile phase of illness again no difference was seen in the infecting viral specific Neut50 titres in patients with either DENV1 or DENV2 infection who proceeded to develop either DF or DHF. Therefore, the Neut50 titres at the beginning of illness, does not appear to associate with subsequent clinical disease severity. Interestingly, in DENV1 infection the Neut50 antibody titres significantly rose in those with DF compared to those with DHF throughout the course of illness, whereas the opposite was seen in DENV2 infection (Neut50 titres rose higher in DHF>DF). This rise in the Neut50 titres throughout the illness was not associated with the viral loads seen in early illness in either DF or DHF patients with DENV1 and DENV2. Therefore, these data show that the degree of viraemia during early illness was not associated with the degree of subsequent rise in Neuto50 antibody titres. In addition, the rise in neutralizing antibody titres does not appear to associate with clinical disease severity but appears to be very different between DENV1 and DENV2. Previous studies have also shown that the degree and duration of viraemia varied between DENV serotypes and there were no differences in the degree of viraemia and kinetics between patients with DF and DHF [35–37]. Collectively, these data show that the rise in the Neut50 antibody titres did not depend on the extent of viraemia; Neut50 titres in the febrile phases in both acute secondary DENV1 and DENV2 were similar in patients who subsequently progressed to develop either DF or DHF.

Although high Neut50 titres are considered to be protective against the infection with the DENV, some individuals with high neutralizing antibodies for a particular DENV serotype later went on to develop DHF when infected with that particular serotype [32]. In this study too, patients who later progressed to develop DHF had similar Neut50 antibody titres as patients who developed DF. Indeed, in a dengue vaccine trial, individuals with high neutralizing antibody titres to certain DENV virus serotypes, were not protected with infection with those serotypes, questioning the Neut50 antibody titres that associate with protection [9, 33]. Therefore, Neut50 antibody titres during the febrile phase measured by *in vitro* assays do not appear to reflect a good correlate of *in vivo* protection.

In summary, the Tfhs appear to significantly expand in acute dengue, which associated with clinical disease severity and plasmablast expansion. This increase in frequency also correlated with DENV-specific IgG, Neut50 antibody titres and with NS1-specific antibody titres. Since evaluation of Neut50 antibody titres in the febrile phase (before patients developed wither DF or DHF) did not appear to associate with clinical disease severity or the degree of viraemia, neutralizing titres alone do not appear to determine protection against severe clinical disease.

## Methods

### Patients for Tfh and plasmablast analysis

40 adult patients with varying severity of acute dengue infection were recruited from the National Institute of Infectious Disease, Sri Lanka following written informed consent. The first blood sample was collected during days 4–8 of illness and a convalescent blood sample was obtained on 21–30 days following the onset of illness. The day on which the patient first developed fever was considered day one of illness. The patients were assessed several times a day and all clinical features such as the blood pressure, urine output, the presence of bleeding manifestations was recorded. Ultrasound scans were performed to determine the presence of fluid leakage in pleural and peritoneal cavities. Full blood counts were performed several times a day throughout the course of illness. Clinical disease severity was classified according to the 2011 World Health Organization (WHO) dengue diagnostic criteria [31]. Accordingly, patients with ultrasound scan evidence of plasma leakage or having a rise in the haematocrit of ≥ 20% from the baseline were classified as having dengue haemorrhagic fever (DHF). Shock was defined as having cold clammy skin, along with a narrowing of pulse pressure of ≤ 20 mmHg. Based on the above criteria, 18 patients were classified as having dengue fever (DF) and 22 patients were classified as having DHF. Twelve healthy individuals were also recruited as healthy controls for the analysis of the frequency and phenotype of Tfh cells and plasmablasts.

### Patients for investigating the changes in Neut50 titres and viral loads throughout the course of illness

In order to investigate the evolution of the Neut50 titres throughout the course of illness and to determine if it associated with clinical disease severity and the degree of viraemia, we recruited a different cohort of 39 patients with acute secondary DENV1 infection (n=22) and with acute secondary DENV2 infection (n=17), Of those with acute secondary DENV1 infection, 12 patients had DF and 10 patients had DHF. Of those with acute secondary DENV2 infection, 8 patients had DF and 9 patients had DHF. They were recruited early during the course of illness, before any of the patients had gone in the vascular leakage (critical phase). Therefore, by obtaining daily blood samples in this cohort of patients, we could study the changes in the Neut50 titres along with the viral loads, well before patients developed any complications. All clinical and laboratory features were recorded as above and clinical disease severity was classified according to the WHO 2011 guidelines [31].

### Ethics statement

Ethical approval was obtained by the Ethics Review Committee of the Faculty of Medical Sciences, University of Sri Jayewardenepura. All patients were recruited following written informed consent.

### Qualitative and quantitative assessment of viral loads

Acute dengue infection was confirmed by quantitative real time PCR and the infecting DENV was serotyped and viral loads were quantified as previously described [38]. Viral RNA was extracted from serum samples using QIAamp Viral RNA Mini Kit (Qiagen, USA) according to the manufacturer’s protocol. Multiplex quantitative real-time PCR was performed as previously described using the CDC real time PCR assay for detection of the DENV and modified to quantify the DENV [39]. Oligonucleotide primers and a dual labelled probe for DENV 1,2,3,4 serotypes were used (Life technologies, USA) based on published sequences. In order to quantify viruses, standard curves of DENV serotypes were generated as previously described by Fernando, S. *et.al*[38].

### Analysis of DENV specific IgM and IgG levels

Dengue antibody assays were performed using a commercial capture-IgM and IgG Enzyme-Linked Immunosorbent Assay (ELISA) (Panbio, Australia). Based on the WHO criteria, patients with an IgM: IgG ratio of >1.2 were classified as having a primary dengue infection, while patients with IgM: IgG ratios <1.2 were categorized under secondary dengue infection[31]. Based on these criteria 13 patients had a primary dengue infection and 27 patients had a secondary dengue infection.

### Flow cytometry for identification of Tfh cells and plasmablasts

Cryopreserved peripheral blood mononuclear cells (PBMC) were thawed in RPMI (Gibco, Life Technologies, USA) at 37°C. Cells were washed in staining buffer (phosphate buffered saline (PBS) containing 2% FBS (Gibco, Life Technologies, USA) and stained with Zombie Green (Biolegend, USA) to determine the percentage of dead cells. For identification and phenotyping of Tfh cells, surface staining was carried out with monoclonal antibodies: anti-CD3 APC/Cy7 (clone OKT3), anti-CD4 pacific blue (clone OKT4), anti-CXCR5 BV711 (clone J252D4), anti-PD-1 BV605 (clone EH12.2H7) and anti-ICOS APC (clone C398.4A), all purchased from Biolegend (San Diego, California, USA). The cells were then fixed in fixation buffer (Biolegend, USA) and permeabilized in intracellular permeabilization wash buffer (Biolegend, USA), followed by staining with anti-IL-21 PE (clone 3A3-N2) (Biolegend, USA) according to the manufacturer’s instructions. Cells were acquired on a Guava easyCyte 12 and analysed with de-novo FCS Express version 6. Tfhs were identified by gating of initially CD3^+^ and CD4^+^ T cells and then gating on CD4^+^ T cells which expressed CXCR5. They were further phenotyped based on their expression of ICOS, PD-1 and IL-21. The gating strategy for phenotyping Tfh cells is shown in supplementary figure 1. Fluorescence Minus One (FMO) controls for each antibody were used to determine the gates.

The following antibodies were used for plasmablast staining: anti-CD19 PE/Cy7 (clone HIB19), anti-CD27 pacific blue (clone M-T271), anti-CD38 APC/Cy7 (clone HIT2) and anti-IgD APC (clone IA6-2), all purchased from Biolegend (San Diego, California, USA). The gating strategy for phenotyping plasmablasts is shown in supplementary figure 2. FMO controls were used to determine the gates for each antibody.

### Determining NS1 antibody levels in patient sera

In order to measure NS1 antibody levels in patient serum samples, an inhouse indirect ELISA was carried out as previously described by Jayathilaka, D. *et.al* [18]. Serum samples diluted at 1:5000 were added to 96-well plates coated with DENV-2 NS1 protein (Native antigen, USA) and blocked with blocking buffer (PBS containing 0.05% Tween 20 and 1 % Bovine serum albumin (BSA). The ELISA was developed with the use of Goat anti-human IgG biotinylated antibody (Mabtech, Sweden), Streptavidin Alkaline Phosphatase enzyme (Abcam, UK) and Para-nitro-phenyl-phosphatase (PNPP) (Thermo Fisher Scientific, USA) substrate. The plate was read on MPSCREEN MR-96A ELISA reader.

### Focus reduction neutralization test (FRNT) for determining neutralizing antibody titres

DENV1 - West Pac 74, DENV-2: S16803 (kindly donated by Prof. Aravinda de Silva) was propagated in C6/36 cell lines and the virus concentration was determined by focus forming assays on Vero-81 cells and expressed as FFU/ml. Using these concentrations, neutralizing antibody titres were assessed by FRNTs. Briefly, Vero-81 cells were seeded overnight on tissue culture treated 96-well plates. An eight-point dilution series of patient serum diluted at 1:5 in DMEM (Gibco, Life Technologies, USA) supplemented with 2% FBS were incubated with a constant amount of DENV1 - West Pac 74 and DENV-2 S16803, for 1 hour at 37 °C with 5% CO_2_. Serum–virus mixtures were then transferred onto the confluent Vero-81 monolayer and incubated for 1 hour at 37 °C. Following the addition of Carboxymethyl Cellulose overlay media, the monolayer was incubated at 37 °C with 5% CO_2_ for 2-3 days. Following incubation, the plates were fixed with 4% Paraformaldehyde (Alfa Aesar, UK) and blocked with a blocking buffer (1X perm buffer (Biolegend, USA) containing 3% Normal Goat Serum) (Sigma, USA). To detect foci a mix of 4G2 and 2H2 (diluted at 1:1000 in blocking buffer) (kindly donated by Prof. Aravinda de Silva) were used as the primary antibodies. HRP conjugated goat anti-mouse IgG (diluted at 1:500 in blocking buffer) (KPL, SeraCare Life Sciences, USA) were used as the secondary antibody and Tru Blue Peroxidase Substrate (KPL, SeraCare Life Sciences, USA) was used as the developing buffer. All assays were done in duplicate.

Spots were counted using RStudio software [40] and data were analysed using GraphPad Prism version 7 software. The neutralization percentages were plotted against the log (1/dilution) values, and the Neut50 was calculated with GraphPad Prism 7.

### Statistical analysis

Data analysis was performed using GraphPad Prism version 7 software. As the data were not distributed normally, differences in means were compared using the Mann-Whitney U test (two-tailed). Wilcoxon signed-rank test was used to compare the differences in means of the acute and convalescent samples. The results were expressed as the median and the interquartile range (IQR). To determine positive and negative correlations, the Spearman’s two-tailed correlation test was used. Statistical significance was set at p<0.05.

## Supporting information

**Supplementary figure 1: The hierarchical gating strategy used to identify Tfhs and to assess their phenotype and functionality.** Cells were first gated on the PBMCS, then the singlets were identified by gating on FSC-height and area, these cells were then gated on the live cells and subsequently on CD3+ cells. The Tfhs were identified as those expressing CD4+ and CXCR5. These Tfhs were subsequently gated on PD-1, ICOS and IL-21 to determine their expression levels.

**Supplementary figure 2: The hierarchical gating strategy used to identify plasmablasts.** Cells were first gated on the PBMCS, then the singlets were identified by gating on FSC-height and area, these cells were then gated on the live cells and subsequently on CD19, CD27 and CD38.

